## Supplemental_figures_S1_S2_S3_S4 for "Contactless micro-elastography of single cells using oscillating microbubbles as shear wave sources"

PNAS

### Supporting Information for
#### Contactless micro-elastography of single cells using oscillating microbubbles as localized shear wave sources

Gabrielle Laloy-Borgna, Maxime Fauconnier, Sibylle Grégoire, Stefan Catheline and Claude Inserra

Gabrielle Laloy-Borgna


**This PDF file includes:**

Figures S1 to S4
Legends for Movies S1 to S3

**Other supporting materials for this manuscript include the following:**

Movies S1 to S3

**Figures**

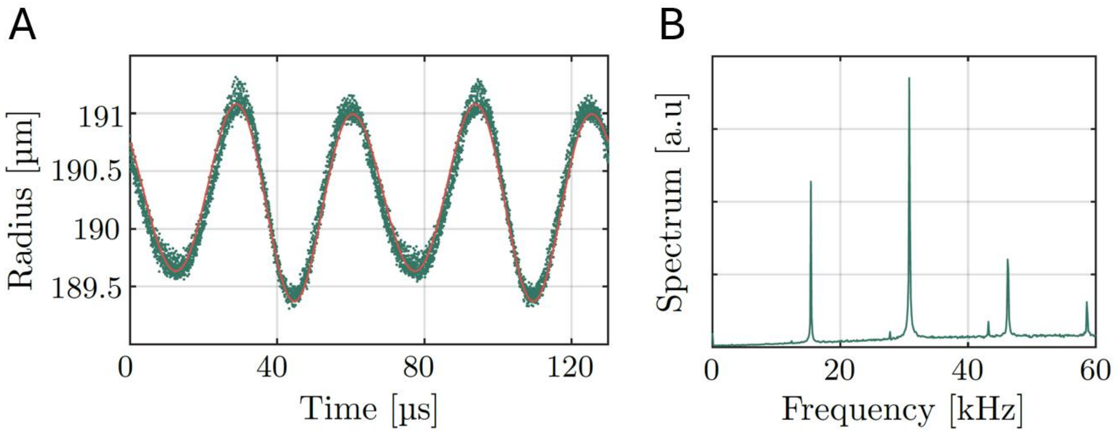


**Fig. S1. Case of a bubble resonant at 15 kHz excited at its resonance frequency. (A) Bubble radius plotted as a function of time. The bubble oscillation is made of a 30 kHz oscillation whose amplitude is modulated by a 15 kHz component. (B) Spectrum of the displacement field measured inside the cell.**

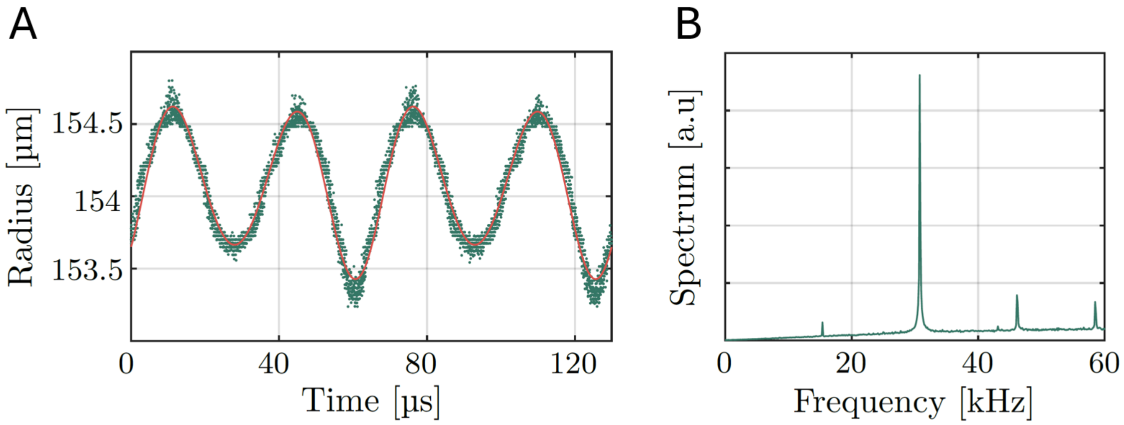


**Fig. S2. Case of a bubble resonant at 30 kHz excited at 15 kHz. (A) Bubble radius plotted as a function of time. The bubble oscillation is made of a 30 kHz oscillation whose amplitude is modulated by a 15 kHz component. (B) Spectrum of the displacement field measured inside the cell.**

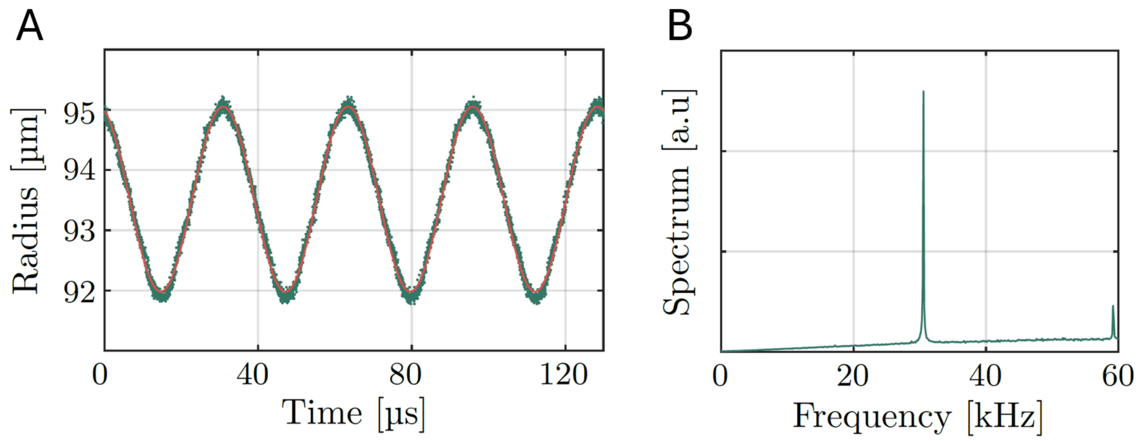


**Fig. S3. Case of a bubble resonant at 30 kHz excited at 15 kHz. (A) Bubble radius plotted as a function of time. The bubble oscillation is made of a pure 30 kHz. (B) Spectrum of the displacement field measured inside the cell.**

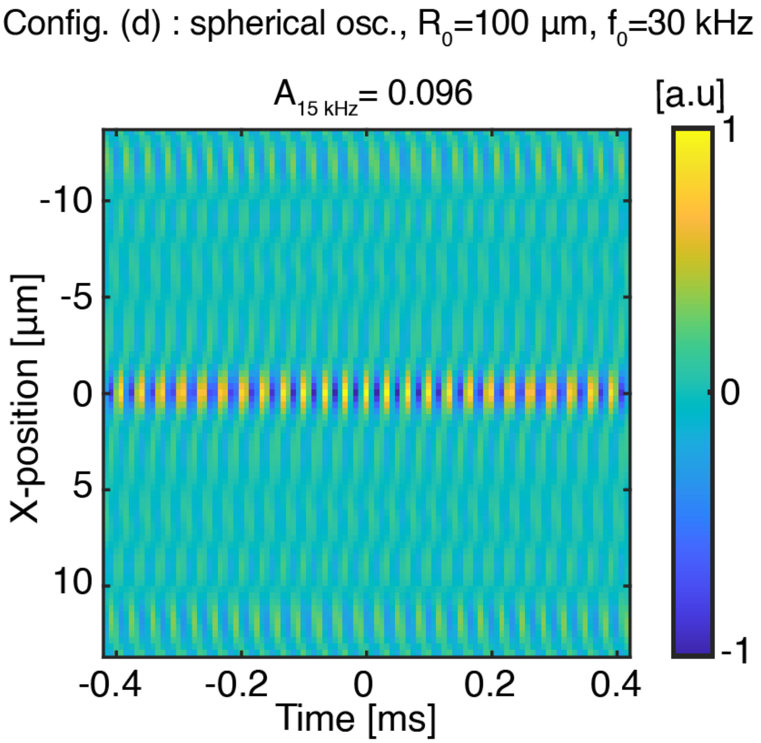


**Fig. S4. Average spatio-temporal correlation of the displacement field measured in the cell for the case of a bubble resonant at 30 kHz excited at 30 kHz. The correlation pattern is made of stationary patterns oscillating over time, but does not show any evidence of wave propagation.**

**Movie S1 (separate file).** Video showing the X-component of the displacement field measured inside the cell, band-pass filtered on the frequency range [14.7 – 16.3] kHz.

**Movie S2 (separate file).** Video showing the X-component of the displacement field measured inside the cell, band-pass filtered on the frequency range [30.5 – 31.5] kHz.

**Movie S3 (separate file).** Video showing the X-component of the displacement field measured inside the cell, band-pass filtered on the frequency range [45.5 - 47] kHz.
